# FusoPortal: An interactive repository of hybrid MinION-sequenced *Fusobacterium* genomes improves gene identification and characterization

**DOI:** 10.1101/305607

**Authors:** Blake E. Sanders, Ariana Umana, Justin A. Lemkul, Daniel J. Slade

## Abstract

Here we present FusoPortal, an interactive repository of *Fusobacterium* genomes that were sequenced using a hybrid MinION long-read sequencing pipeline, followed by assembly and annotation using a diverse portfolio of predominantly open-source software. Significant efforts were made to provide genomic and bioinformatic data as downloadable files, including raw sequencing reads, genome maps, gene annotations, protein functional analysis and classifications, and a custom BLAST server for FusoPortal genomes. FusoPortal has been initiated with eight complete genomes, of which seven were previously only drafts that varied from 24-67 contigs. We showcase that genomes in FusoPortal provide accurate open reading frame annotations, and have corrected a number of large genes (>3 kb) that were previously misannotated due to contig boundaries. In summary, FusoPortal (http://fusoportal.org) is the first database of MinION sequenced and completely assembled *Fusobacterium* genomes, and this central *Fusobacterium* genomic and bioinformatic resource will aid the scientific community in developing a deeper understanding of how this human pathogen contributes to an array of diseases including periodontitis and colorectal cancer.

**Importance:** In this study, we report a hybrid MinION whole genome sequencing pipeline, and describe the genomic characteristics of the first eight strains deposited in the FusoPortal database. This collection of highly accurate and complete genomes drastically improves upon previous multi-contig assemblies by correcting and newly identifying a significant number of open reading frames. We believe this resource will result in the discovery of proteins and molecular mechanisms used by an oral pathogen, with the potential to further our understanding of how *F. nucleatum* contributes to a repertoire of diseases including periodontitis, pre-term birth, and colorectal cancer

## Introduction

Multiple *Fusobacterium* species are oral pathogens that can infect a broad range of human organ and tissue niches.^1,2^ *Fusobacterium nucleatum* has recently been connected with colorectal cancer (CRC),^3,4^ with studies showing this bacterium induces a pro-inflammatory microenvironment and chemoresistance against drugs used to treat CRC.^5–7^ Despite the importance of *Fusobacterium* in human diseases, a lack of complete genomes of biomedically relevant isolates has hindered protein cataloging and virulence factor identification. Many *Fusobacterium* draft genomes have been sequenced and partially assembled using short-read technologies (454 Life Sciences), leaving complete genome assembly difficult due to repeat regions. The reference genome of *F. nucleatum* subsp. *nucleatum* ATCC 25586 was completed using cosmid and λ phage technologies to achieve long reads (10-35 kb) and whole genome assembly.^8^ However, we show in a parallel study that while this genome was assembled into one complete chromosome, we uncovered a 452 kb inversion using our sequencing and assembly methods. With the emergence of next generation long-read sequencing (Pacific Biosciences, Oxford Nanopore MinION), assembling whole genomes has become standard and affordable for academic research settings. The recent combination of MinION long-read and Illumina short-read technologies to scaffold and polish DNA sequencing data, respectively, has created a robust pipeline for bacterial genome completion and subsequent gene identification and characterization.^9,10^

The motivation for complete sequencing and assembly of *Fusobacterium* genomes came from our discovery that bioinformatic analysis identified a high percentage of large genes (~3,000-12,000 bp) in the *F. nucleatum* 23726 genome that appeared to be missing critical protein domains at either the N- or C-terminus (e.g. >2000 amino acid deletions). We show that these genomic discrepancies are not isolated to *F. nucleatum* 23726 by correcting a substantial number of large proteins (up to 5300 amino acid corrections) in all eight genomes. Since the largest proteins in *Fusobacterium* are autotransporters of the Type 5 secretion system, these new genomes will be important to reevaluate the virulence factor landscape of these pathogenic bacteria.

To provide ease of use and data accessibility to the community, we have used this study to launch the FusoPortal repository, which provides the first eight completely sequenced, assembled, and annotated *Fusobacterium* genomes using MinION and Illumina technology. While databases including KEGG, NCBI, and Uniprot are crucial for researchers to find open reading frames, our goal was to create a central database in which researchers interested in *Fusobacterium* biology could obtain high-quality data in an easy to navigate platform. The FusoPortal repository framework has been developed to allow additional genomes to efficiently be added, with a goal of assembling 25 previously incomplete genomes spanning a broad range of *Fusobacterium* species. In summary, genomes and bioinformatic analysis available in the FusoPortal repository provides key resources to further determine how this understudied pathogen contributes to a variety of human infections and diseases. Herein we highlight not only how users can interact with the FusoPortal website, but additional bioinformatic analysis that was made possible due to improved genome sequencing and assembly.

## Results and Discussion

### Genome sequencing, assembly, and annotation

In this study, we have successfully completed seven new *Fusobacterium* genomes, and have resequenced the widely referenced strain *F. nucleatum* subsp. *nucleatum* 25586. Each genome now consists of a single circular chromosome, and in the case of *F. varium* 27725, we report the discovery of a 42 kb circular plasmid that contains 70 protein genes ranging from 38 to 1678 amino acids. For each genome, we have provided access to all protein and RNA encoding genes, as well as highlighting the previously unidentified CRISPR systems. In Figure 1, we describe the workflow used to complete genome sequencing, assembly, annotation, functional prediction, and implementation of the interactive FusoPortal repository. We made a concerted effort to use open-source software when possible, to make this workflow reproducible and accessible to the scientific community.

**Figure 1:**
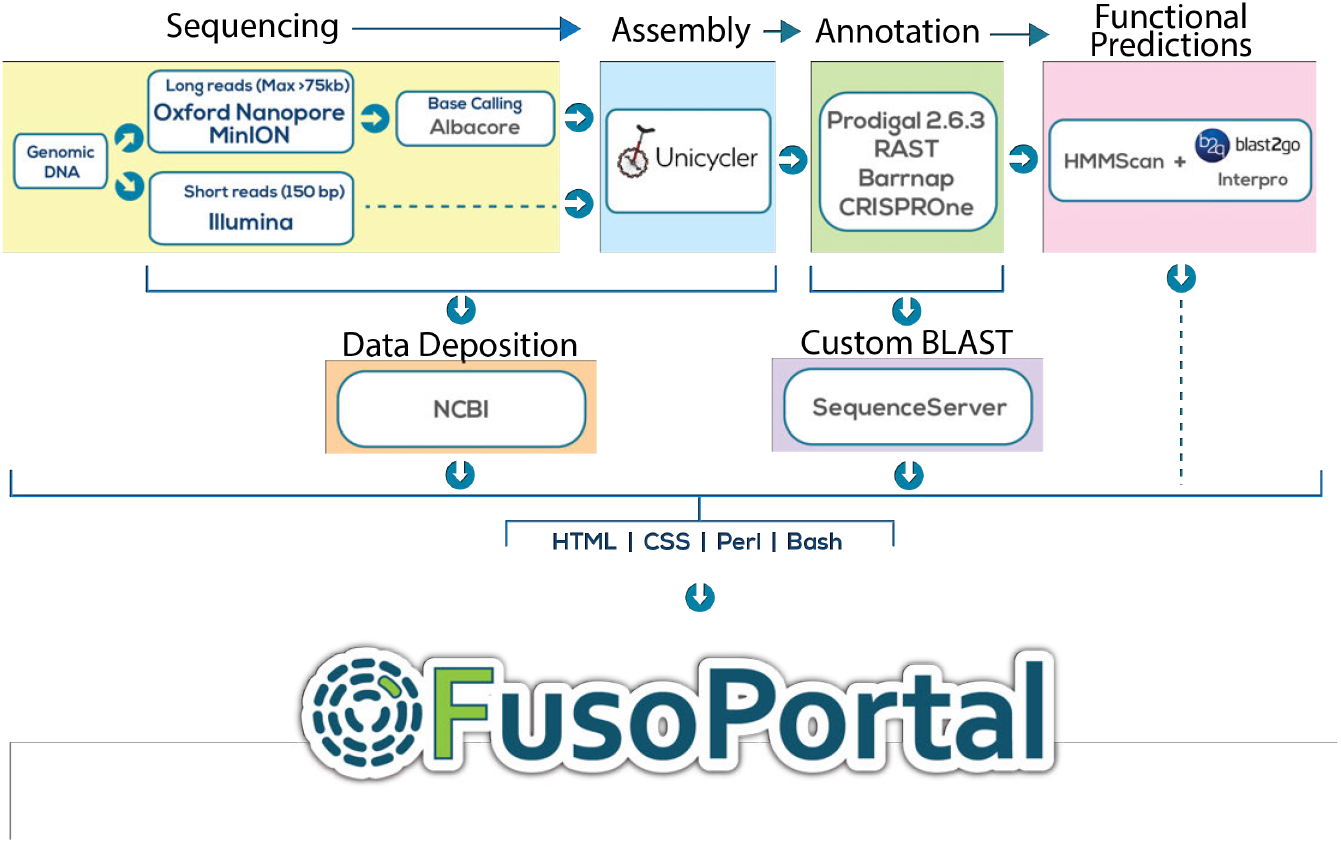
Schematic of all genomic, bioinformatic, and scripting workflows to create FusoPortal.

We highlight that the largest change in gene number from a reference genome to our single chromosome build was *F. necrophorum* 1_1_36S, which was previously reported with 3197 protein encoding genes. We analyze our new prediction of 2125 genes, and show that this number is much more consistent with the genome size when comparing all *Fusobacterium* genomes (Figure 2A,Table 1). These data show that the previous annotation contained an abundance of short open reading frames, and the complete genome increases the average gene length by more than 250 bp; in which the average gene size of ~1000 bp agrees well with the remaining seven *Fusobacterium* genomes (Figure 2B).

**Table 1.** Statistics and NCBI accession numbers for FusoPortal genomes.

| Species | Strain | NCBI Genbank Sequence | NCBI Contigs<sup>§</sup> | Size (mb) | Protein genes | FusoPortal Genbank Sequence | FusoPortal Contigs | Size | Protein genes |
| --- | --- | --- | --- | --- | --- | --- | --- | --- | --- |
| *F. nucleatum* | 23726 | GCF_000178895.1 | 67 | 2.23 | 1983 | GCA_003019785.1 | 1 | 2.30 | 2111 |
| *F. nucleatum* | 25586 | GCA_000007325.1 | 1 | 2.17 | 1968 | GCA_003019295.1 | 1 | 2.18 | 2019 |
| *F. varium* | 27725 | GCA_000159915.2 | 39 | 3.30 | 3008 | GCA_003019655.1 | 1 | 3.35 | 3054<sup>Δ</sup> |
| *F. ulcerans* | 49185 | GCA_000158315.2 | 49 | 3.49 | 3191 | GCA_003019675.1 | 1 | 3.54 | 3230 |
| *F. mortiferum* | 9817 | GCA_000158195.2 | 44 | 2.67 | 2538 | GCA_003019315.1 | 1 | 2.72 | 2631 |
| *F. gonidiaformans* | 25563 | GCA_000158835.2 | 24 | 1.70 | 1618 | GCA_003019695.1 | 1 | 1.68 | 1617 |
| *F. periodonticum* | 2_1_31 | GCA_000158215.3 | 61 | 2.55 | 2375 | GCA_003019755.1 | 1 | 2.54 | 2388 |
| *F. necrophorum* | 1_1_36S | GCA_000242215.1 | 40 | 2.31 | 3197 | GCA_003019715.1 | 1 | 2.29 | 2125 |

<sup>§</sup> Contig - Continuous stretch of genomic DNA with no breaks.
<sup>Δ</sup> Includes 70 genes from 42 kb plasmid: GCA\_003019655.1

^$^ Contig - Continuous stretch of genomic DNA with no breaks.

^Δ^ Includes 70 genes from 42 kb plasmid: GCA_003019655.1

### Correcting large, protein-encoding reading frames

As expected, the largest change in the *F. necrophorum* 1_1_36S genome is in the number of genes that are now annotated with 1000 amino acids or more. In fact, the combined analysis of all eight genomes shows that we have corrected a significant percentage of proteins that were previously annotated with <1000 amino acids and now contain >1000 amino acids, or proteins that were already >1000 amino acids and now contain expanded protein open reading frames (Figure 2C). The most extreme example was a previously annotated protein in the *F. necrophorum* 1_1_36S genome that was 960 amino acids (EHO18576.1), and has now been annotated as a 6248 amino acid protein in FusoPortal. *F. mortiferum* 9817 did not have any amino acid additions in proteins over 1000 residues, but we note that the largest protein in this genome is 1602 residues (EEO35225.2), and as expected, proteins in this range have fewer open reading frame errors than larger proteins frequently found in *Fusobacterium* genomes. We highlight that in the biomedical significant and genetically tractable strain *F. nucleatum* 23726, there were several large stretches of amino acids added to proteins that are homologous to the previously characterized virulence protein Fap2.^11–13^ In addition, we correct an abnormally large number of genes in the reference strain *F. nucleatum* 25586, which we attribute to improved bacterial annotation software, and not genomic errors in the previous complete genome.

**Figure 2:**
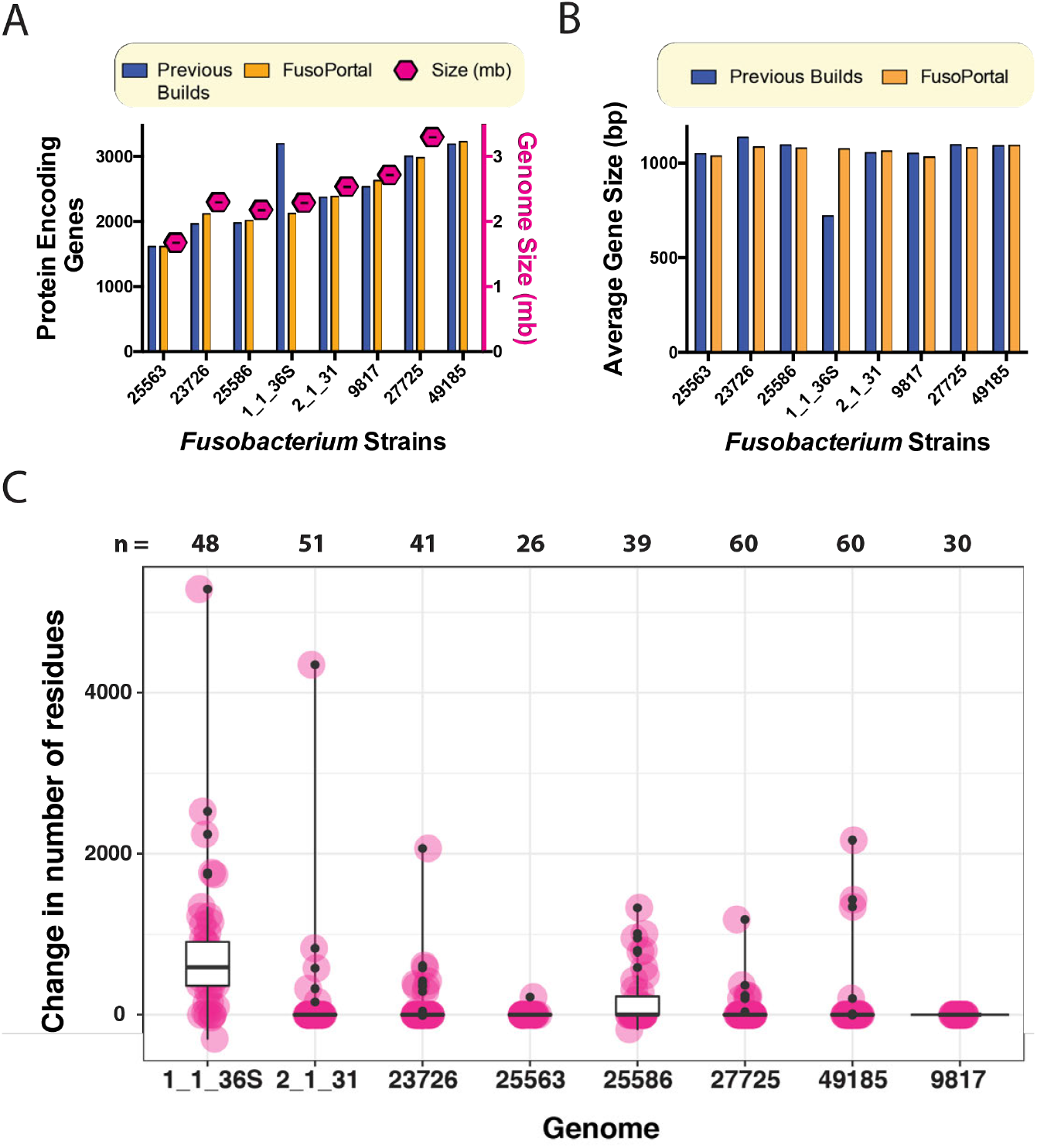
Correction of open reading frames in *Fusobacterium* genomes. **(A)** A comparison of the genome size to the number of protein encoding genes per genome in both the NCBI and FusoPortal genomes. **(B)** A comparison of all proteins for average gene size in NCBI and FusoPortal genomes. **(C)** Analysis of all proteins 1000 residues and above in annotated FusoPortal proteomes, and how many residues were added (or in rare cases removed) compared to the previous annotations present in the NCBI database. n = number of proteins 1000 residues or greater in each genome.

### Whole genome phylogenetic analysis of a diverse group of *Fusobacterium*

With newly identified genes in our reported *Fusobacterium* genomes, we set out to create a phylogenetic tree to compare to previous reports. By using all open reading frames from each genome to build a tree, we show that while *F. nucleatum* 23726 and *F. nucleatum* 25586 are quite close phylogenetically, the remaining genomes are quite diverse in their genetic makeup Figure 3. Because of the diversity of these eight genomes, there are a large number of unique gene clusters, which has been analyzed previously and was hypothesized to govern differences in intracellular invasion and virulence potential.^14^

**Figure 3:**
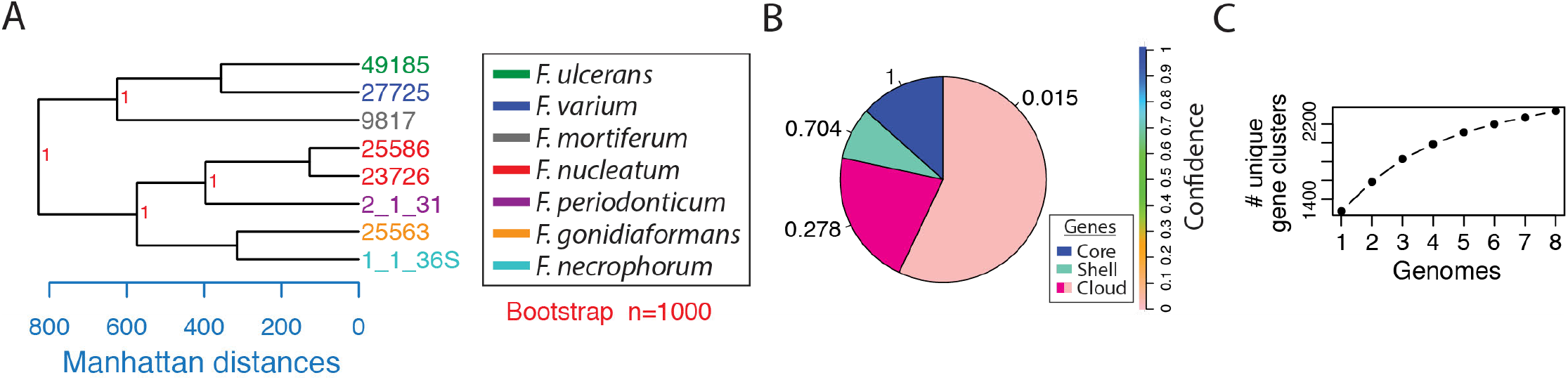
Phylogenetic analysis of complete *Fusobacterium* genomes. **(A)** Tree created using complete proteomes in the microPan package in R. Bootstrap values are indicated in red. **(B)** micropan analysis of pan-genome gene families with models predicting the percentage of core genes, shell genes (present in most genomes), and cloud genes (found in few genomes). **(C)** Analysis of the number of unique gene clusters found among the eight complete FusoPortal genomes.

### Features of the FusoPortal repository

FusoPortal was built on an HTML5 framework, and therefore is functional on all full sized computers and mobile devices. The home page gives a description of features available in FusoPortal, and provides links to all genomes and bioinformatic data. Links to all raw (Illumina and MinION DNA reads) and processed genomes and annotations are available as show in Figure 4A. In Figure 4B, we highlight a genome map in which directional arrows represent clickable open reading frames that send the user to individual pages containing DNA and protein coding sequences in FASTA format. We have also analyzed all proteins on FusoPortal using HMMer^15^ (downloadable.out files), and have produced custom linked Interpro pages with full bioinformatic analysis (Figure 4C). These resources alleviate the need for users to exit the site to acquire bioinformatic data, and provide functional predictions for entire proteomes.

**Figure 4:**
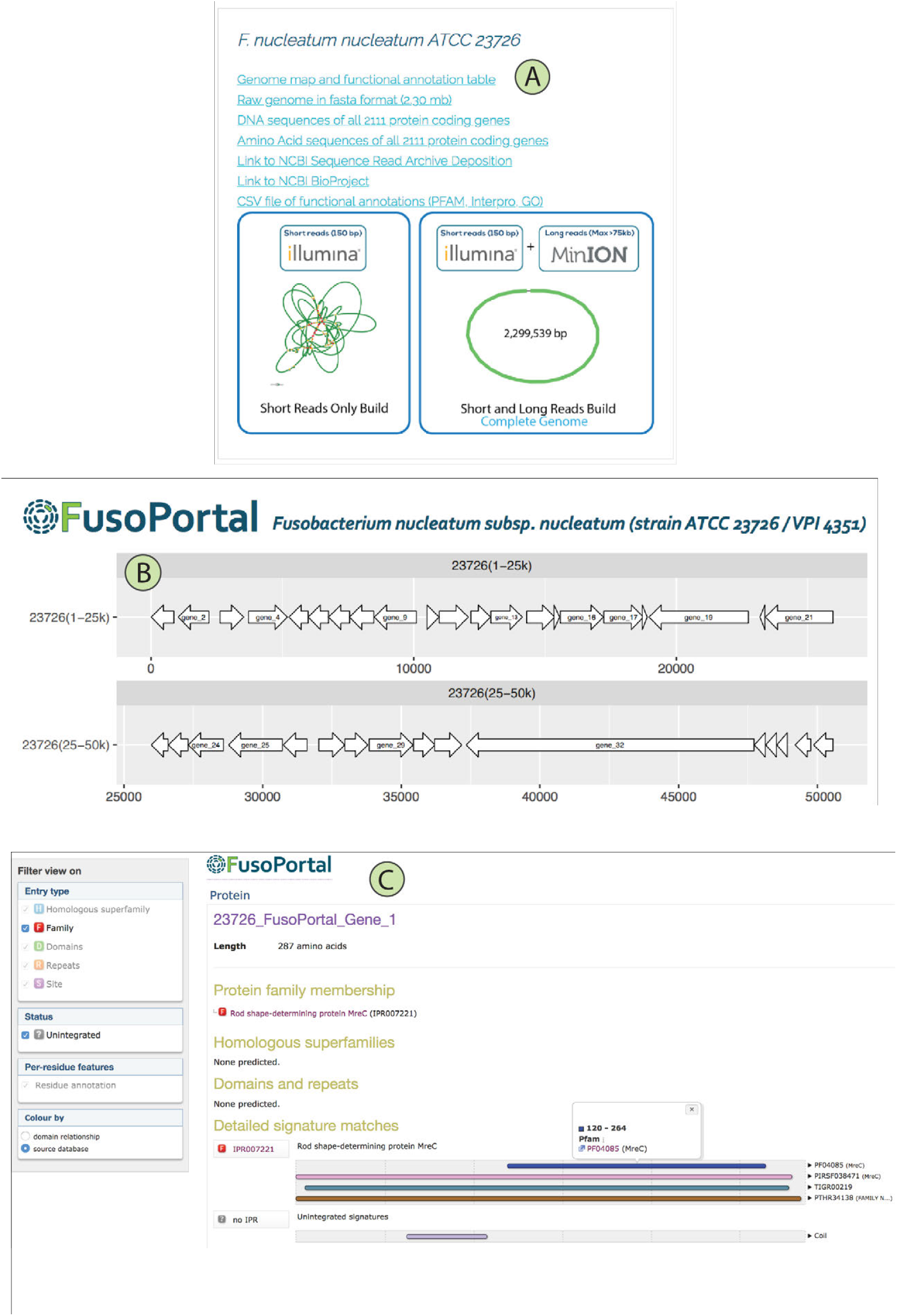
Navigating the FusoPortal web interface for genome analysis. **(A)** Links are provided for downloading all genomic data (raw and analyzed), with Bandage plots to show the completeness of each genome build. **(B)** Whole genome maps were produced with links to custom web pages for each gene that provide access to all genomic and bioinformatic analysis. **(C)** Each gene contains a full Interpro analysis page with links to multiple functional prediction databases (e.g. PFAM, TIGR, Gene3D, CDD, GO).

### A custom BLAST database to search FusoPortal

To additionally aid in virulence factor identification, we built a custom BLAST server using the open source software Sequenceserver (www.sequenceserver.com).^16^ All eight genomes, including the *F. varium* 27725 plasmid, can be searched using a DNA or protein sequence input as shown in Figure 5A-B. Results are provided as alignments with e-values (customizable inputs for thresholds), and all acquired sequences and alignments can be downloaded in various formats (Figure 5C-D).

**Figure 5:**
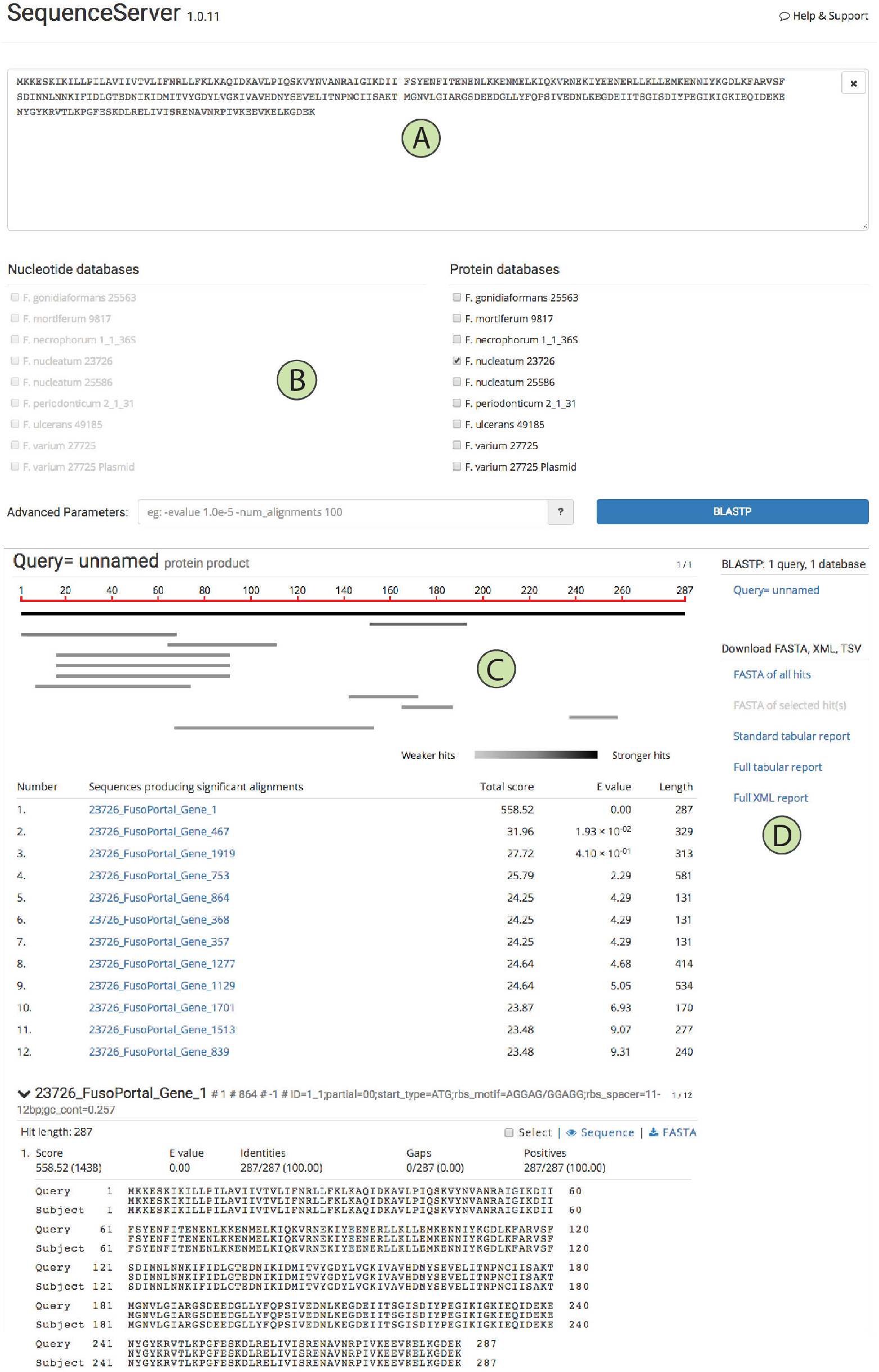
A custom BLAST database to search FusoPortal genomes built with SequenceServer. (A) Data entry panel that auto-detects protein or DNA sequences. (B) Genomes available that users can choose to search. (C) BLAST results for identified genes. (D) Links to downloadable files.

### Conclusion

In conclusion, FusoPortal is a database of fully sequenced, annotated, and bioinformatically characterized *Fusobacterium* genomes that provides a central location to increase our understanding of these virulent bacteria contribute to a wide range of diseases. We aim to expand this resource to provide additional value to the scientific community, which will allow for more detailed comparative genomics for virulence prediction of *Fusobacterium* species.

## Materials and Methods

### Data to populate FusoPortal

Detailed sequencing statistics and assembly methods are reported in a concurrent publication by Slade and colleagues.^17^ Briefly, genomic DNA was isolated from cultured *Fusobacterium* and sequenced using MinION (Oxford Nanopore Technologies) and Illumina platforms. Genomes were assembled using the open source software package Unicycler^10^. Gene annotations were obtained using Prodigal,^18^ RAST,^19^ Barrnap,^20^ and CRISPRone^21^ and prediction of protein function was achieved with the stand alone HMMer program^15^ and InterPro^22^ from within the Blast2GO software platform^23^. A graphical representation of methods for DNA sequencing, genome assembly, bioinformatics, and construction of FusoPortal are presented in Figure 1.

### Development of interactive genome maps

Protein open reading frames predicted by Prodigal were used to identify boundaries that were used in the R package gggenes to create complete genome maps. For each genome, DNA gene coordinates were placed in a CSV file and imported into the gggenes package in Studio to create linear genome maps. Maps were then manually loaded into Adobe Illustrator for custom FusoPortal formatting, followed by adding links to genes using Adobe Acrobat Pro. Final genome maps were exported as PDF files for incorporation into the FusoPortal website. All CSV files with gene coordinates and a template R file is provided on our Open Science Framework database http://osf.io/2c8pv.

### Construction of FusoPortal using HTML and automated field filling scripts

FusoPortal was built using HTML5 and CSS, and automation of HTML page filling with genomic and bioinformatic data was implemented using custom Bash and Perl scripts. All scripts are available on an Open Science Framework database run by Daniel J. Slade at http://osf.io/2c8pv. Scripts were developed to automate the acquisition of custom HMMER models for all genomes, produce linear genome maps using gggenes in the Studio package. For populating FusoPortal HTML pages, scripts were created to extract Prodigal gene coordinates from.gbk files, extract gene and protein sequences, and populate web pages with Interpro functional annotations from.csv files for population into HTML pages. Interpro analysis pages were produced by Blast2GO, and the FusoPortal logo was added to each page through scripting.

### Construction of a FusoPortal custom BLAST server

The FusoPortal custom BLAST server was built using the Sequenceserver software package.^16^ Briefly, a custom Apache server was implemented on an Amazon Light Sail private server and all FusoPortal genomes containing protein open reading frames in DNA (.fna) and amino acid (.faa) formats were implemented as guided by the Sequenceserver advanced setup and configuration documentation. This BLAST server can be accessed at http://18.216.121.101/blast/.

### Accession numbers

All accession numbers for data is provided in Table 1.

## Acknowledgements

This work was supported by the USDA National Institute of Food and Agriculture and startup funding from the Department of Biochemistry at Virginia Tech to Daniel J. Slade.

## Author Contributions

**Blake E. Sanders**, Data curation, Code writing, Writing-review and editing; **Ariana Umana**, Data curation, Code writing, Writing-review and editing; **Justin A. Lemkul**, Code writing, Writing-review and editing; **Daniel J. Slade**, Conceptualization, Data curation, Formal analysis, Supervision, Funding acquisition, Validation, Methodology, Writing-original draft, Project administration, Writing-review and editing.

## Author ORCIDs

**Blake E. Sanders**: https://orcid.org/0000-0002-5841-122X

**Ariana Umana**: https://orcid.org/0000-0002-1941-8656

**Justin A. Lemkul**: https://orcid.org/0000-0001-6661-8653

**Daniel J. Slade**: https://orcid.org/0000-0001-5634-7220

